# A modified BCG with depletion of enzymes associated with peptidoglycan amidation induces enhanced protection against tuberculosis in mice

**DOI:** 10.1101/2023.05.03.539199

**Authors:** Moagi T. Shaku, Peter Um, Karl L. Ocius, Alexis J. Apostolos, Marcos M. Pires, William R. Bishai, Bavesh D. Kana

## Abstract

Mechanisms by which *Mycobacterium tuberculosis* (Mtb) evades pathogen recognition receptor activation during infection may offer insights for the development of improved tuberculosis (TB) vaccines. Whilst Mtb elicits NOD-2 activation through host recognition of its peptidoglycan-derived muramyl dipeptide (MDP), it masks the endogenous NOD-1 ligand through amidation of glutamate at the second position in peptidoglycan sidechains. As the current BCG vaccine is derived from pathogenic mycobacteria, a similar situation prevails. To alleviate this masking ability and to potentially improve efficacy of the BCG vaccine, we used CRISPRi to inhibit expression of the essential enzyme pair, MurT-GatD, implicated in amidation of peptidoglycan sidechains. We demonstrate that depletion of these enzymes results in reduced growth, cell wall defects, increased susceptibility to antibiotics and altered spatial localization of new peptidoglycan. In cell culture experiments, training of monocytes with this recombinant BCG yielded improved control of Mtb growth. In the murine model of TB infection, we demonstrate that depletion of MurT-GatD in BCG, resulting in unmasking of the D-glutamate diaminopimelate (iE-DAP) NOD-1 ligand, yields superior prevention of TB disease compared to the standard BCG vaccine. This work demonstrates the feasibility of gene regulation platforms such as CRISPRi to alter antigen presentation in BCG in a bespoke manner that tunes immunity towards more effective protection against TB disease.

## Introduction

Tuberculosis (TB) caused by *Mycobacterium tuberculosis* (Mtb) remains a leading cause of death from an infectious disease worldwide (WHO, 2022). Despite the availability of the Bacille Calmette Guerin (BCG) TB vaccine, approximately 2 billion people worldwide are latently infected with Mtb and represent a reservoir of future active disease (Trunz et al., 2006). BCG is the only licensed TB vaccine and has been in use since the 1920s with close to 100 million infants vaccinated worldwide (Trunz et al., 2006). BCG protects against TB meningitis and miliary TB in children, but lacks efficacy against pulmonary TB in adults (Martinez et al., 2022); hence, improved TB vaccines remain an urgent public health priority.

Innate immune pattern recognition receptors (PRRs) have evolved to sense unique pathogen-associated molecular patterns (PAMPs) that are often essential components of infecting organisms (Li and Wu, 2021). Bacterial peptidoglycan (PG) is one such PAMP, and it is detected by the PRRs NOD-1, which recognizes the D-isoglutamate diaminopimelate (iE-DAP) segment of PG, and NOD-2 which detects the related muramyl dipeptide (MDP) portion of PG (Li and Wu, 2021). Activation of NOD-1 triggers the production of pro-inflammatory cytokines through nuclear factor κB (NF-κB) and mitogen-activated protein kinase (MAPK) pathways and similarly, NOD-2 activation leads to upregulation of NF-κB activity (Caruso et al., 2014).

While many gram-negative pathogens express abundant levels of the NOD-1 ligand iE-DAP (Caruso et al., 2014), pathogenic mycobacteria including Mtb, *M. bovis*, and the *M. bovis*-derived BCG strains possess an immune subversion system which enzymatically masks NOD-1 antigenic structure and thereby enables escape from NOD-1 mediated immune containment (Maitra et al., 2019). The enzyme pair encoded by the *murT* (Mb3739 in *M. bovis*)-*gatD* (Mb3749) operon forms a glutaminase and an amidotransferase complex, which amidates iE-DAP to form iQ-DAP, thus avoiding NOD-1 detection (**Figure 1a** and **Figure S1**). As amidation of D-isoglutamate to D-isoglutamine during PG maturation in mycobacteria is required for subsequent PG cross-linking, MurT and GatD are important for mycobacterial cell wall integrity and genetic screens have confirmed their essentiality for *in vitro* survival (de Wet et al., 2020;Maitra et al., 2021).

**Figure 1:**
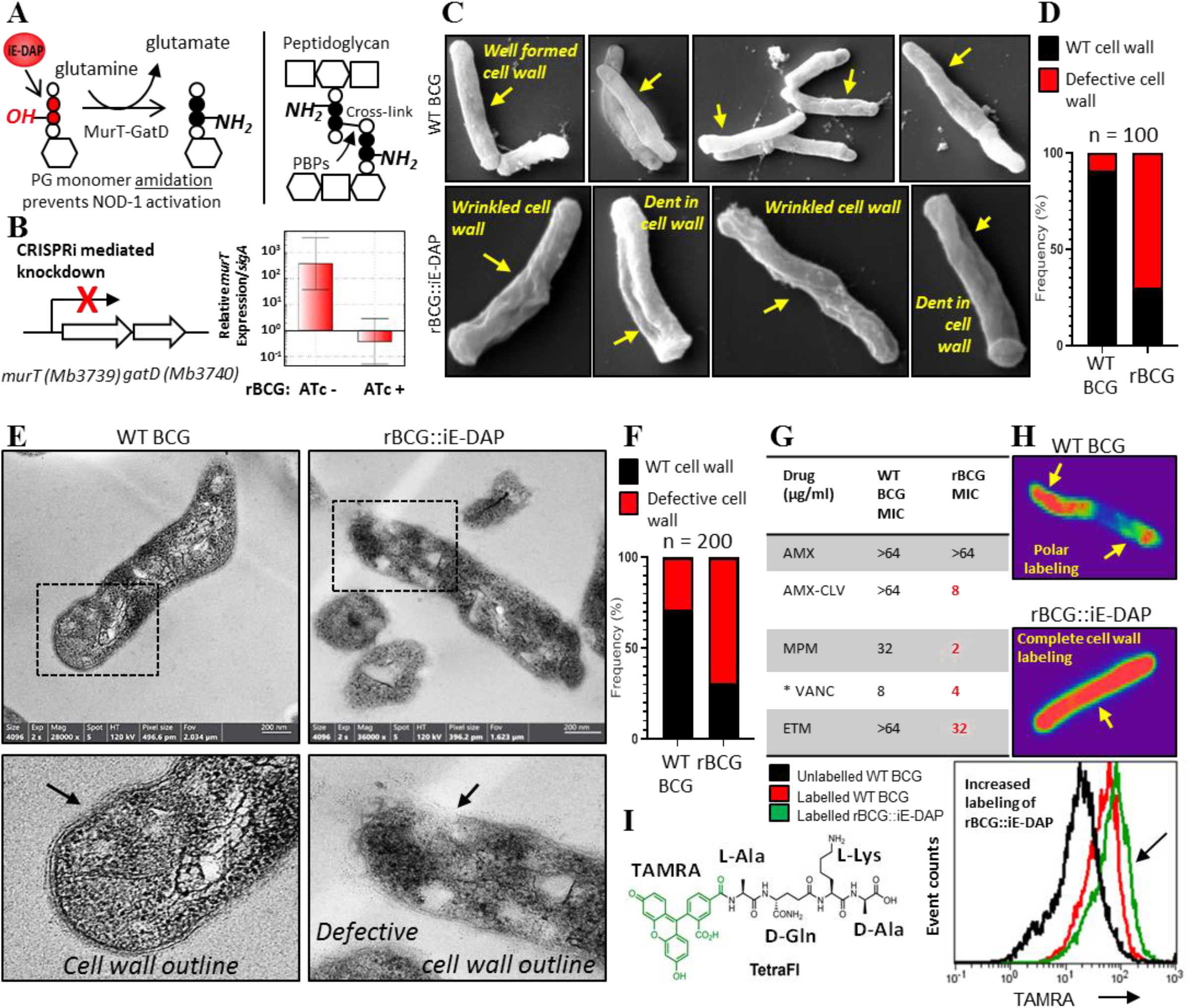
Phenotypic characterization of rBCG::iE-DAP. (A) Schematic representation of MurT-GatD mediated G precursor amidation. (B) qPCR data for CRISPRi depletion of MurT-GatD in rBCG::iE-DAP. CRISPRi tivation with ATc (200 ng/ml) resulted in inhibition of *murT* expression. (C) Scanning electron micrographs of WT CG ((n=45 micrographs, 100 cells counted) and rBCG::iE-DAP grown in media supplemented with 200 ng/ml ATc. epletion of MurT-GatD in rBCG::iE-DAP (n=48 micrographs, 100 cells counted) causes outer cell wall defects rinkled and dents in the cell wall). (D) Frequency of cells with cell wall defects. (E) Transmission electron crographs of WT BCG and rBCG::iE-DAP grown in media supplemented with 200 ng/ml ATc. Depletion of MurT-atD causes cell wall defects. (F) Frequency of cells with cell wall defects. (G) Minimum inhibitory concentration of tibiotics targeting cell wall biosynthesis against WT BCG and rBCG::iE-DAP grown in media supplemented with 0 ng/ml Atc. AMX: Amoxicillin, CLV: Clavulanate, MPM: Meropenem, VANC: Vancomycin and ETM: hionamide. Depletion of MurT-GatD in rBCG::iE-DAP causes increased sensitivity to cell wall targeting tibioics. (H) MurT-GatD depleted cells labelled with fluorescent BODIPY-FL vancomycin reveals side-wall beling. (I) Flow cytometry analysis of WT BCG and rBCG::iE-DAP cells labelled with a PG amidation reporter obe (TAMRA-L-Ala-D-glutamine-L-Lys-D-Ala [TetraFI]). Depletion of MurT-GatD causes increased labeling th the PG amidation reporter probe indicative of reduced PG amidation in the MurT-GatD depleted cells.

We hypothesized that depletion of MurT-GatD in BCG would result in increased abundance of the NOD-1 ligand (iE-DAP), thus enabling enhanced immunogenicity of the recombinant vaccine strain. We used a CRISPRi platform for targeted inhibition of transcription of the amidotransferase complex - MurT-GatD essential for PG amidation in mycobacteria (i.e. modification of iE-DAP to iQ-DAP) to develop a recombinant BCG vaccine (rBCG::iE-DAP) engineered to activate NOD-1 during vaccination. Our results show that MurT-GatD levels can be conditionally depleted in BCG without complete loss of viability and that compared to the wildtype (WT) BCG, vaccination of mice with the MurT-GatD-depleted rBCG gives superior containment of Mtb proliferation in lungs.

## Results

### Construction of rBCG::iE-DAP

Using the CRISPRi gene expression knockdown system we generated a derivative of plasmid pLRJ965 (Rock et al., 2017) that conditionally expresses dCas9 from *Streptococcus thermophiles* and a 17 base short guide RNA (sgRNA) sequence that target the *murT-gatD* operon upon exposure to anhydrotetracycline (ATc) or doxycycline (Dox) to create plasmid PLRJ965+*murT*sgRNA (**Figure S2a and S2b**). This plasmid was introduced into BCG-Pasteur to generate a recombinant BCG strain called rBCG::iE-DAP. We showed that following ATc induction, the relative mRNA levels of the full length *murT* transcript were 1000-fold lower in rBCG::iE-DAP when compared with the uninduced rBCG strain (**Figure 1b)**.

Next, we evaluated the impact of MurT-GatD depletion on BCG viability. CRISPRi mediated inhibition of *murT*-*gatD* transcription in rBCG::iE-DAP by supplementation of growth media with an increasing concentration of ATc [0-500 ng/ml] resulted in growth inhibition of the recombinant strain (**Figure S2c**). This is consistent with earlier knockdown of a MurT homologue in *Mycobacterium smegmatis* (MSMEG_6276) which revealed a growth defect upon CRISPRi mediated MSMEG_6276 depletion (de Wet et al., 2020).

### MurT-GatD depletion in BCG causes cell wall defects

To determine the effects of MurT-GatD depletion on BCG cell-wall structure, we performed scanning and transmission electron microscopy (SEM, TEM). As shown in (**Figure 1c**), SEM revealed a well-formed typical mycobacterial outer cell wall structure in WT BCG whereas rBCG::iE-DAP cells displayed a wrinkled outer-cell wall structure, sometimes with indentations indicative of altered cell wall biosynthesis upon MurT-GatD depletion. Quantification of SEM fields revealed a 70% increase in the frequency of bacilli with these defects in rBCG::iE-DAP relative to the WT BCG (**Figure 1d**). Consistent with this, TEM revealed a typical multi-layered mycobacterial cell wall outline, with visible layers in WT BCG in comparison to the defective cell wall structure in rBCG::iE-DAP, without a clear cell wall outline as shown in **Figure 1e and Figure S3**, (Paul and Beveridge, 1993;Brennan, 2003). Upon counting individual cells in TEM fields, we observed a 65% increase in the frequency of wall defects in rBCG::iE-DAP compared with the WT BCG strain (**Figure 1f**).

We hypothesized that reduced PG cross-linking due to MurT-GatD depletion, and the concomitant cell wall defects, might enable antibiotics which target cell wall biosynthesis to be more potent in rBCG::iE-DAP compared to WT BCG. Indeed, as shown in **Figure 1g**, MurT-GatD knockdown was associated with a 2-to 16-fold decrease in the minimal inhibitory concentrations of the recombinant strain for amoxicillin-clavulanate, meropenem, vancomycin, and ethionamide, each of which targets either PG biosynthesis or PG-dependent accessory glycolipids. To further confirm the reduced levels of PG cross-linking, we stained MurT-GatD depleted rBCG::iE-DAP cells with BODIPY-FL vancomycin—a fluorescent probe which specifically labels uncrosslinked PG. As shown in the confocal fluorescence micrographs in **Figure 1h**, BODIPY-FL vancomycin displayed complete cell wall labeling of the rBCG::iE-DAP cells, in contrast, only the poles of WT BCG cells were labeled. This corresponds to the known polar elongation of BCG cells and the relative abundance of new, uncross-linked PG at the cell poles (Aldridge et al., 2012;Joyce et al., 2012).

To specifically demonstrate that MurT-GatD depletion resulted in reduced amidation of iE-DAP, we used a fluorogenic amidated, synthetic tetrapeptide, TetraFl (TAMRA fluorophore-L-Ala-D-Gln-L-Lys-D-Ala). This amidated, D-Gln-containing probe is incorporated into mycobacterial PG by the activity of PG cross-linking L,D-transpeptidases which require the amidation modification on one of the PG stem peptides to form the cross-link (Pidgeon et al., 2019). The delayed or deficient cross-linking in the cell wall due to MurT-GatD depletion led us to speculate that more of the amidated probe will be incorporated into existing PG. As seen in **Figure 1i**, labeling of the MurT-GatD depleted cells showed a greater incorporation of the tetrapeptide fluorophore than in WT BCG, thus confirming that rBCG::iE-DAP displayed greater exposure of the iE-DAP, NOD-1 antigenic structure.

### rBCG::iE-DAP is responsive to anhydrotetracycline activation and causes increased TNFα expression in bone marrow derived macrophages (BMDMs)

To test the hypothesis that inhibition of MurT-GatD expression in rBCG::iE-DAP enhances the immunogenicity of the recombinant strain through increased expression of the NOD-1 ligand iE-DAP, we first infected IFNγ-activated BMDMs with rBCG::iE-DAP and supplemented the growth media with increasing concentrations of ATc to assess activation of the CRISPRi system *ex vivo* and also to compare growth to WT BCG infected cells. The growth of the strains was recorded by plating for colony forming unit (CFU) counts at day 3 and day 5 post-infection. At day 3 bacterial containment was observed for all strains but was most prominent for rBCG::iE-DAP strains treated with ATc. Dose dependent inhibition of growth of rBCG::iE-DAP was observed at day 5, with 500 ng/ml ATc (the maximum concentration used) resulting in a ∼3 fold difference in growth inhibition of rBCG::iE-DAP in comparison to WT BCG and rBCG::iE-DAP without ATc supplementation, (**Figure 2a**). Secondly, we performed ELISA experiments to assess the expression of the pro-inflammatory cytokine TNFα as rBCG::iE-DAP is designed to express the NOD-1 ligand iE-DAP and potentially induce increased NF-κB activation leading to a potentially increased pro-inflammatory response. Activation of rBCG::iE-DAP by supplementation of growth media with ATc resulted in a dose dependent increase in TNFα expression in comparison to WT BCG in IFNγ-activated BMDMs, although this was statistically insignificant between strains, while TNFα expression remained low for both WT BCG and rBCG::iE-DAP strains when used for infection of unactivated BMDMs (**Figure S4**). These results demonstrate that rBCG::iE-DAP is responsive to activation *ex vivo* and signals for increased pro-inflammatory cytokine expression.

**Figure 2:**
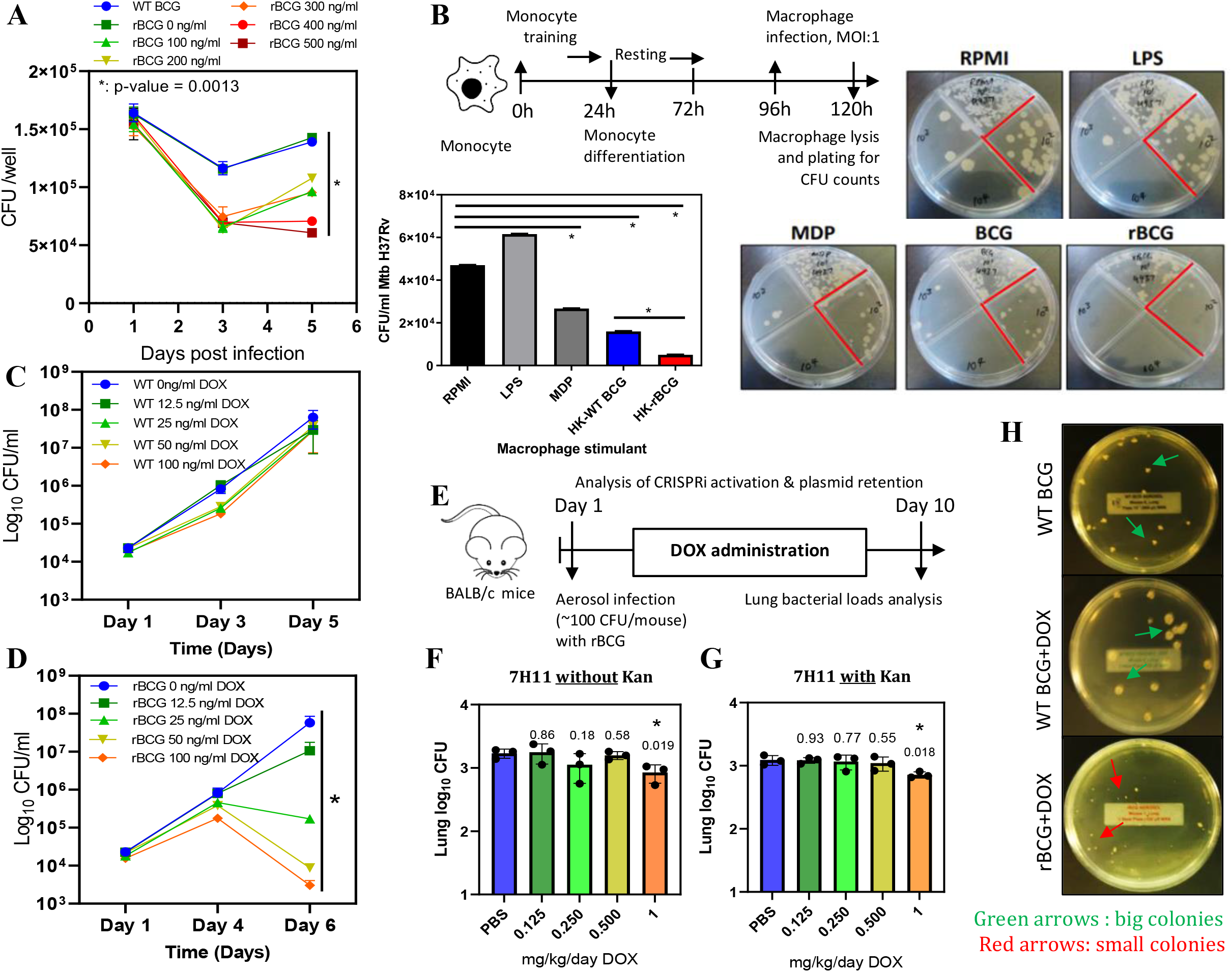
Survival of rBCG::iE-DAP in IFNγ activated bone marrow derived macrophages (BMDMs), training monocytes and activation with doxycycline. (A) IFNγ activated BMDMs (1 × 10^6^ cells) were culture media for induction of the CRISPRi system in CG::iE-DAP at concentrations ranging from 100 ng/ml – 500 ng/ml and growth of the strains was assessed after 3 d 5 days. Supplementation of culture media with 400-500 ng/ml ATc resulted in significantly reduced growth of CG::iE-DAP on days 3 and 5 in comparison to WT BCG. (B) Training of U937 monocytes with heat-killed (HK)-CG::iE-DAP compared to HK-WT BCG. Shown is also the representative CFU plates for the experiment. acrophages derived from HK-rBCG::iE-DAP trained monocytes control Mtb growth better than HK-WT BCG ined macrophages. (C-D) CFU counts of *in vitro* grown WT BCG by (C) and of rBCG::iE-DAP (D) grown in complete 7H9 medium at varying concentrations of Dox. (E) Determination of the Dox concentration for activation rBCG::iE-DAP *in vivo*. Mice were aerosol infected with ∼2.5 log_10_ CFU of rBCG::iE-DAP and Dox (0.125-1 /kg/day) - was administered by oral gavage for 10 days. (F-G) CFU counts from the experiment shown in panel E. dose of 1 mg/kg/day Dox resulted in reduced growth of rBCG::iE-DAP indicative of sufficient activation of the ISPRi system. Lung homogenates were plated on both 7H11 with (G) and without (F) kanamycin (25 µg/ml) to ess the loss of the CRISPRi plasmid during *in vivo* growth. There was no difference in the growth of recovered bacteria on media containing kanamycin, indicating the presence of the CRISPRi plasmid in recovered bacteria. P-ues are given above the graphs. (H) Plates showing the colony size of rBCG::iE-DAP+Dox compared to WT BCG WT BCG+Dox, recovered from the lungs of aerosol infected mice. student *t*-test was used for statistical analysis. p-value <0.05.

### rBCG::iE-DAP *in vitro* trained macrophages control Mtb H37Rv growth

WT BCG trains macrophages in a NOD-2 dependent manner and as a result, killing of Mtb is enhanced if the trained cells are exposed to Mtb at a later stage (Kleinnijenhuis et al., 2012;Kaufmann et al., 2018). We hypothesized that rBCG::iE-DAP engineered to express the NOD-1 ligand upon activation with ATc will lead to enhanced macrophage training activity, resulting in better control of Mtb growth compared to WT BCG trained macrophages. To test this we used an *in vitro* macrophage training assay to assess the Mtb killing ability of rBCG::iE-DAP trained macrophages. Lipopolysaccharide (LPS) and murein dipeptide (MDP) were used as controls and a cells-only (RPMI) control was also included. LPS activates toll like receptor (TLR)-4 leading to monocyte activation (Fujihara et al., 2003) and MDP activates NOD-2 leading to macrophage training (van der Heijden et al., 2018). As shown in **Figure 2b**, heat-killed rBCG::iE-DAP trained macrophages displayed increased control of Mtb H37Rv compared to heat-killed WT BCG trained macrophages and MDP trained macrophage. Macrophages derived from LPS stimulated monocytes did not control Mtb growth. Based on these promising findings, we proceeded to test rBCG::iE-DAP in the murine model of TB infection.

### rBCG::iE-DAP activation *in vitro* and in mice-aerosol infections with doxycycline

Doxycycline (Dox), a tetracycline analog, is used in *in vivo* TB models for temporal regulation of mycobacterial gene expression (Miow et al., 2021). The CRISPRi platform used for generation of rBCG::iE-DAP is also based on a Dox-responsive TetR-*tetO* unit which in the presence of doxycycline leads to expression of the CRISPRi system and subsequent transcriptional inhibition of *murT*-*gatD* (Rock et al., 2017). To assess the activation of rBCG::iE-DAP with Dox, the strain was grown in an increasing range of Dox concentrations to assess the activation of CRISPRi *in vitro*, a WT BCG+Dox control experiment was also included. Activation of CRISPRi in rBCG::iE-DAP with Dox resulted in a dose dependent reduction of rBCG::iE-DAP growth (**Figure 2c-d)**, which was corroborated when growth was assessed by CFU counts while WT BCG was not affected by Dox supplementation (**Figure 2c-d**).

To test the activation of rBCG::iE-DAP *in vivo* and to determine the minimum effective dose of Dox, we aerosol infected BALB/c mice with ∼100 CFU of rBCG::iE-DAP and administered Dox for 10 days at doses ranging from 0.125 - 1 mg/kg/day by oral gavage (**Figure 2e**). Administration of 1 mg/kg/day resulted in a significant reduction in growth of rBCG::iE-DAP in the lungs of the mice (**Figure 2f**). To assess retention of the CRISPRi plasmid (PLRJ965+*murT*sgRNA, which has a kanamycin [Kan] resistance cassette) by rBCG::iE-DAP *in vivo*, we plated lung homogenates also on media containing Kan and found that recovered rBCG::iE-DAP bacilli formed similar CFU counts on media with or without Kan (**Figure 2f-g**). As shown in **Figure 2h**, at 8 weeks post infection, rBCG::iE-DAP bacilli recovered from the lungs of infected mice formed small colonies on solid agar in comparison to recovered WT BCG bacilli, indicative of the long term efficacy of 1 mg/kg/day Dox *in vivo* for CRISPRi activation. These results demonstrate retention of the CRISPRi plasmid by rBCG::iE-DAP *in vivo*.

### rBCG::iE-DAP induces enhanced protection against *Mycobacterium tuberculosis* infection in mice compared to WT BCG

To assess the protective efficacy of rBCG::iE-DAP against TB infection relative to the standard BCG vaccine, we immunized groups of BALB/c mice (n=5 per group) intradermally with WT BCG or rBCG::iE-DAP (**Figure 3a**). rBCG::iE-DAP immunized mice received a Dox dose by oral gavage at 1 mg/kg/day for activation of CRISPRi *in vivo* and we also included a Saline+Dox group, a WT-BCG+Dox group and a rBCG::iE-DAP without Dox group as controls for the vaccination experiment. The immunized mice receiving Dox were weighed weekly for 6 weeks prior to Mtb challenge to assess the effect of daily Dox administration on the health of the mice (**Figure 3a)**. We assessed the percentage weight change of mice receiving Dox relative to the no-Dox groups and found that the weights of the different groups remained within 80-100% of baseline with few significant differences (**Figure 3b**).

**Figure 3:**
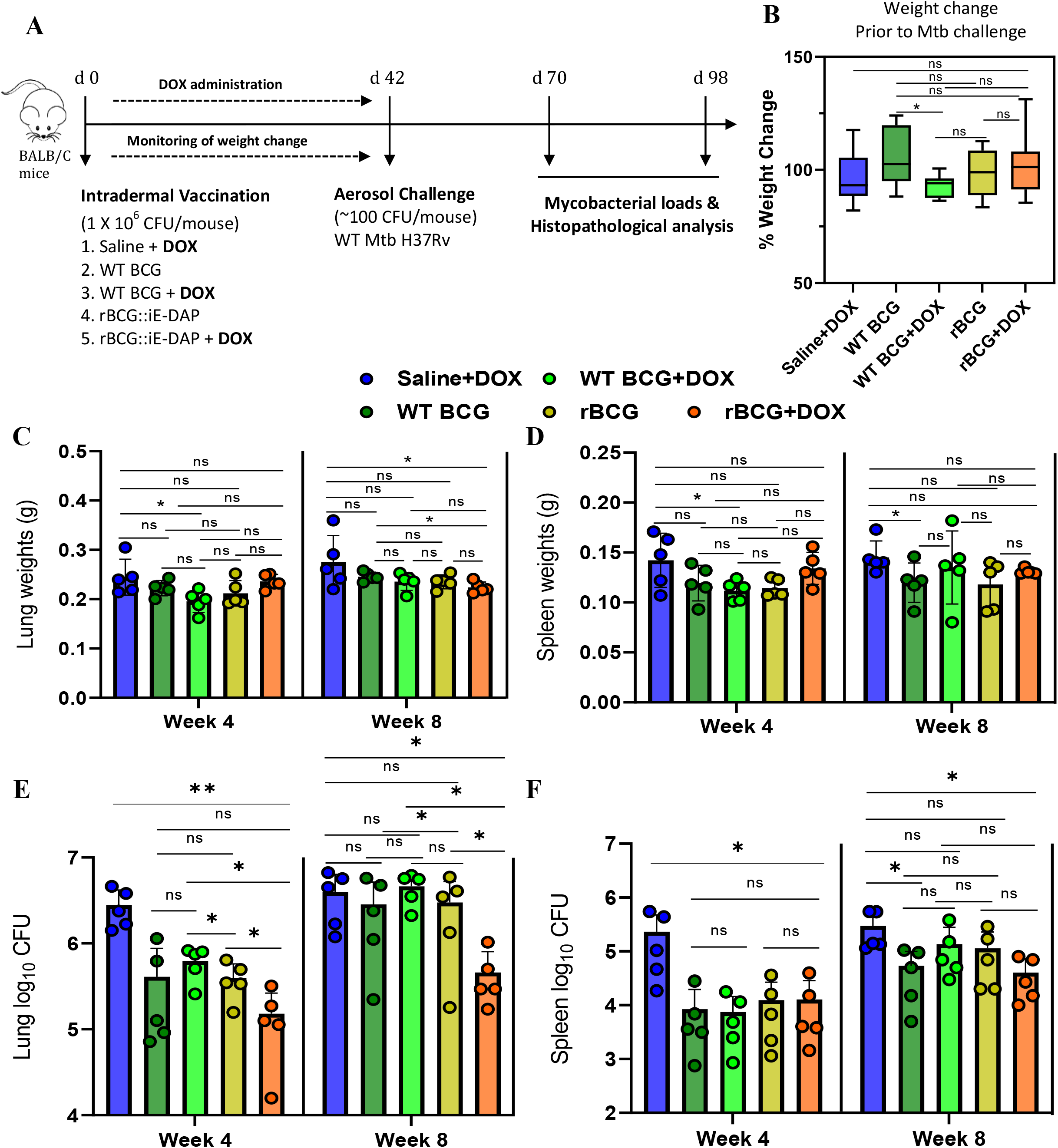
Efficacy of rBCG::iE-DAP in comparison to standard WT BCG for protection against Mtb H37Rv fection in mice. (A) Schematic representation of the mouse immunization and Mtb H37Rv challenge protocol. (B) Percentage weight change at week 6 (d42) immediately prior to Mtb challenge. (C-D) Lung and Spleen weights week 4 and week 8 post-challenge with Mtb. (E-F) Lung and Spleen bacterial burdens at week 4 and week 8 st-challenge with Mtb. rBCG::iE-DAP+Dox was superior to WT BCG or WT BCG+Dox in controlling Mtb 37Rv growth in the lungs at week 4 and week 8 post challenge with Mtb. * :p-value <0.05.

After 6 weeks, the immunized mice were challenged with ∼100 CFU of Mtb H37Rv via the aerosol route and mycobacterial loads were determined in lungs and spleens at 4 and 8 weeks post challenge (**Figure 3a)**. At 4 weeks post Mtb challenge mice were sacrificed to assess lung pathology and bacterial burden in the lungs and spleens. As seen in **Figure 3c**, the WT BCG+Dox group displayed lower lung weights, the WT BCG and rBCG::iE-DAP both without Dox-treatment groups displayed similar lung weights while the Saline+Dox and rBCG::iE-DAP+Dox groups displayed increased lung weights indicative of increased lung inflammation. The Saline+Dox group displayed increased spleen weights compared to the other groups (**Figure 3d**). As shown in **Figure 3e and 3f**, respectively, analysis of lung and spleen bacterial burden at 4 weeks post infection revealed that rBCG::iE-DAP+Dox was superior to WT BCG and WT BCG+Dox in protecting against Mtb challenge in the lungs and reduced bacterial dissemination to the spleen similar to WT BCG or WT BCG+Dox. At 8 weeks post Mtb challenge, the rBCG::iE-DAP+Dox group displayed reduced lung weights indicative of control of bacterial burden and indeed, analysis of lung bacterial burden corroborated findings at 4 weeks that rBCG::iE-DAP+Dox was superior to WT BCG or WT BCG+Dox in controlling Mtb growth in the lung (**Figure 3c and 3e**). At week 8, WT BCG or WT BCG+Dox vaccination both displayed waning efficacy in this model, as previously shown (Henao-Tamayo et al., 2015;Dwivedi et al., 2022). In the spleen, rBCG::iE-DAP+Dox displayed similar efficacy to WT BCG or WT BCG+Dox for control of infection compared to the Saline+Dox group (**Figure 3f**).

### Histopathological analysis of lung pathology after vaccination with rBCG::iE-DAP compared to WT BCG post Mtb challenge

As shown in **Figure 4a-b**, histopathological analysis of haematoxylin and eosin (H&E) stained lung samples from the vaccinated and Mtb challenged mice indicated that rBCG::iE-DAP+Dox immunized mice presented with early increased lung inflammation compared to WT BCG+Dox vaccinated mice. At 8 weeks post infection also, rBCG::iE-DAP+Dox immunized mice presented with increased inflamed sections of lung area compared to WT BCG+Dox immunized mice suggestive of sustained inflammation for control of infection (**Figure 4c-d**). The increased early inflammation in rBCG::iE-DAP+Dox immunized mice is reflective of early induction of anti-tuberculous immune responses, which were able to control growth early before establishment of infection and the sustained inflammation at 8 weeks post challenge is suggestive of enhanced immune responses during chronic disease stage which enable control of disease progression as shown in **Figure 3e**.

**Figure 4:**
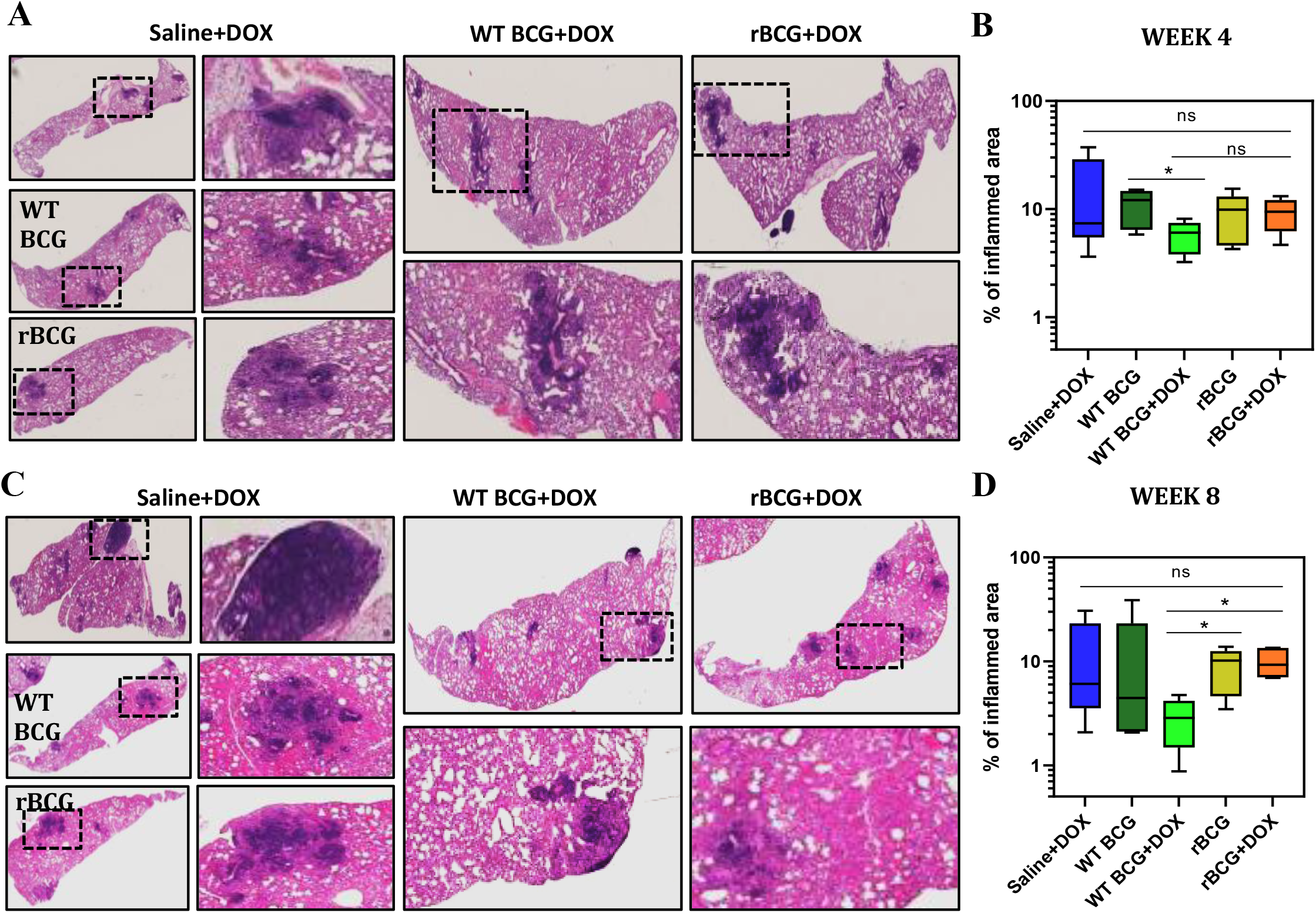
Histopathological analysis of lung samples. (A) Histological Haematoxylin and Eosin (H&E) staining of ng samples at week 4 post Mtb challenge. (B) Analysis of percentage of inflamed area (indicated with black arrows) from each mouse lung per immunized group (n=5 per group), shows that rBCG::iE-DAP+Dox immunized ice present with early lung inflammation compared to WT BCG+Dox. (C) Histological Haematoxylin and Eosin H&E) staining of lung samples at week 8 post Mtb H37Rv infection. (D) Analysis of percentage of inflamed area from each mouse lung (n=5 per group), shows that rBCG::iE-DAP+Dox immunized mice present with sustained lung inflammation compared to WT BCG and WT BCG+Dox. The percentage inflamed area was evaluated using ImageJ software (NIH) and plotted as whisker boxplots (whiskers represent minimum and maximum values) and a student *t*-test was used for statistical analysis. * :p-value <0.05.

## Discussion

The BCG vaccine has been in use for decades at birth and is effective at preventing TB in young children. However, BCG does not provide protection against TB infection in adults and has failed to eradicate the TB pandemic despite it being the only licensed TB vaccine (Dockrell and Butkeviciute, 2022). This has spurred the need to develop novel TB vaccine candidates with varied modes of action to replace BCG or boost BCG, which still remains the gold-standard for next generation TB vaccine development (Martinez et al., 2022). BCG possesses a number of immune evasion mechanisms similar to those used by Mtb during infection to avoid immune killing that can be targeted for deletion to enhance its efficacy. For example, rBCG strains further attenuated by deletion of immune evasion genes such as *sapM* (Festjens et al., 2019), *nuoG* (Gengenbacher et al., 2016) or *zmp1* (Sander et al., 2015) among others, have been developed and these show enhanced immunogenicity and efficacy against Mtb infection in animal models.

Immune evasion genes that are also essential for BCG viability are attractive targets to be studied for development of next generation rBCGs with enhanced efficacy. For example, genes encoding essential enzymes involved in the biosynthesis of potent immune-modulating cell wall lipids such as trehalose dimycolate (TDM), di- and tri-acylglycerols, pthiocerol dimycocerosates (PDIMs) and phenolic glycolipids (PGLs) are potential targets of gene regulation platforms to study their role in limiting BCG efficacy. Selective chemical removal of these lipids from BCG (i.e. delipidation of BCG) has shown the ability to increase BCG efficacy in preventing Mtb infection in mice (Moliva et al., 2019). Gene regulation platforms, including CRISPRi are ideal platforms to study the effect of such essential immunomodulatory BCG enzymes that can be targeted to enhance BCG efficacy. Indeed, recently a CRISPRi based rBCG (rBCG::CRISPRi-AftC) designed for the truncation of the anti-inflammatory cell wall associated lipoglycan – lipoarabinomannan (LAM) into the pro-inflammatory lipomannan derivative (LM) upon CRISPRi mediated depletion of the enzyme arabinofuranosyltransferase C (AftC, required for addition of D-arabinan branches on LM) was shown to enhance the immunogenicity of BCG by upregulating the expression of TNFα, a major pro-inflammatory cytokine (Madduri et al., 2022).

We targeted essential genes (*murT*-*gatD*) required for PG amidation in BCG with the recently developed CRISPRi platform (Rock et al., 2017) to create rBCG::iE-DAP designed to activate the NOD-1 pathway by expression of iE-DAP. Phenotypic characterization of rBCG::iE-DAP post CRISPRi activation, with SEM and TEM displayed changes in the outer cell wall surface and structure of the cell wall in rBCG::iE-DAP respectively, consistent with the essentiality of MurT-GatD during PG biosynthesis. These defects were also associated with increased sensitivity to cell wall targeting antibiotics (amoxicillin+clavulanate, meropenem, vancomycin and ethionamide), confirming the essentiality of MurT-GatD mediated amidation of PG fragments for cell wall biosynthesis as previously described in mycobacteria and other bacterial species (Münch et al., 2012;Liu et al., 2017;Maitra et al., 2021). As PG amidation by the MurT-GatD complex is required for PG cross-linking by penicillin binding proteins and L,D-transpeptidases in mycobacteria (Ngadjeua et al., 2018), we further analysed the level of PG cross-linking in rBCG::iE-DAP by labeling the cells with BODIPY-FL vancomycin, a fluorescent vancomycin derivative binding uncross-linked nascent PG monomers (Miao et al., 2020). We found that transcriptional repression of MurT-GatD expression in rBCG::iE-DAP was associated with complete cell wall labeling with this probe indicative of reduced PG cross-linking in rBCG::iE-DAP, due to lack of MurT-GatD enzymatic activity. To probe for the reduction of PG amidation in rBCG::iE-DAP, we used a previously developed amidation reporter probe, TetraFl (Pidgeon et al., 2019) to label MurT-GatD depleted cells and this showed increased labeling in rBCG::iE-DAP, indicative of reduced amidation upon transcriptional repression of MurT-GatD expression.

We then first tested rBCG::iE-DAP in an *in vitro* monocyte training assay (Bekkering et al., 2016) to assess the efficacy of this strain in training innate responses of macrophages. BCG induces a NOD-2 dependent trained immunity in monocytes resulting in epigenetic and metabolic reprogramming of monocytes which differentiate into macrophages with increased bactericidal properties (Kleinnijenhuis et al., 2012;Blok et al., 2015). We therefore tested rBCG::iE-DAP trained macrophages for their *in vitro* Mtb killing ability and found that in contrast to WT BCG trained macrophages, rBCG::iE-DAP trained macrophages displayed enhanced Mtb killing ability as measured by CFU counts 24 hours post infection.

We further show that rBCG::iE-DAP is responsive to activation in BMDMs using ATc and also in a mouse aerosol infection model using a minimal Dox dose, as Dox was previously shown to have immunomodulatory effects (Miow et al., 2021). Activation of rBCG::iE-DAP in BMDMs resulted in increased TNFα expression measured by ELISA providing evidence that expression of iE-DAP in rBCG::iE-DAP enhances immunogenecity of BCG. These results bolstered our enthusiasm for investigation of rBCG::iE-DAP *in vivo* as a potential TB vaccine candidate.

We demonstrated that intradermal vaccination of mice with rBCG::iE-DAP followed by administration of Dox for 6 weeks for CRISPRi mediated repression of MurT-GatD expression resulted in superior protection from Mtb challenge in the lungs of the immunized mice compared to the WT BCG vaccine. Our vaccination experiments also included a WT BCG+Dox control group to rule out the role of Dox mediated immunomodulatory effects in enhancing WT BCG vaccine efficacy post Mtb challenge when compared with rBCG::iE-DAP+Dox efficacy. Administration of Dox to WT BCG vaccinated mice (i.e. WT BCG+Dox group) did not enhance WT BCG vaccine efficacy against Mtb challenge when compared to the WT BCG without Dox control group, while rBCG::iE-DAP+Dox shows increased protection at both 4 and 8 weeks post Mtb challenge. We also observed a waning efficacy of the WT BCG vaccine at week 8 post Mtb infection, which has been previously reported (Henao-Tamayo et al., 2015;Dwivedi et al., 2022), however, interestingly vaccination with rBCG::iE-DAP+Dox remained effective at this time point.

Histopathological analysis of lung sections from the immunized and Mtb challenged mice shows that rBCG::iE-DAP+Dox induces early immune infiltration to the lung compared to WT BCG+Dox and this is maintained until 8 weeks post infection, providing early and sustained protection against Mtb challenge. The immune correlates of protection induced by rBCG::iE-DAP are the subject of our future studies. As we used a CRISPRi based knockdown strategy to create rBCG::iE-DAP, the next step would be to generate suppressor mutants of rBCG::iE-DAP that allow complete deletion of the *murT*-*gatD* operon in BCG to generate a strain that is not based on CRISPRi as a TB vaccine candidate. Collectively, our work demonstrates that MurT-GatD can be targeted to develop a new TB vaccine candidate.

## Materials and Methods

### Bacterial strains and Culture conditions

#### Growth conditions for *E. coli* DH5α and derivative strains

*E. coli* DH5α and derivative strains were grown in Luria-Bertani broth (LB) or on Luria-Bertani agar (LA) at 37°C with supplementation of the media with appropriate antibiotics. The antibiotic concentration used was as follows: Kanamycin (Kan): 50 µg/ml. Liquid cultures were grown at 37°C with shaking at a 100 rpm.

#### Growth conditions for Mycobacterial and derivative strains

*M. bovis* BCG, *M tuberculosis* H37Rv and the recombinant BCG::CRISPRi MurT-GatD strain were grown at 37°C in Middlebrook 7H9 broth supplemented with OADC enrichment, 0.5% glycerol, 0.05% Tween 80 and appropriate antibiotics (hereafter referred to as Middlebrook 7H9 broth) or on Middlebrook 7H11 agar supplemented with OADC enrichment and 0.5% glycerol and appropriate antibiotics. The antibiotic concentration used for kanamycin was 50 µg/ml.

#### Construction of rBCG::iE-DAP

The programmable mycobacterial CRISPRi system for repression of gene transcription was used as previously described by Rock et al. 2017, to generate the recombinant BCG::CRISPRi strain – rBCG::iE-DAP (14). Briefly, the CRISPRi system utilizes a catalytically-inactivated *anhydrotetracycline*/doxycycline (ATc/Dox)-inducible CRISPRi dcas9 from *Streptococcus thermophiles*, which is directed by a (ATc/Dox)-inducible short-guide RNA (sgRNA) to specific target genes to prevent transcription initiation or elongation (Rock et al., 2017). sgRNAs were designed with the CRISPRi sgRNA design tool - https://pebble.rockefeller.edu/. The sgRNA sequence (top and bottom oligos) were annealed and cloned into BsmBI-digested CRISPRi vector PLJR965 (**Supplementary Figure 2a-b**). These plasmids were introduced into *M. bovis* BCG by electroporation.

#### Quantitative real-time PCR (qPCR)

qPCR was performed using Sso Fast Evagreen Supermix (BioRad) as per manufacturer’sinstructions. Briefly, 20 µl reactions were set up, each containing 10 µl Evagreen Supermix, 0.75 µl forward primer (Mb3739Fwd: gtcaaacgattcggtcagctg) (10 µM), 0.75 µl reverse primer (Mb3739Rev: gattcaccgagcctggcag) (10 µM), 2 µl cDNA and nucleasefree water. All reactions were incubated in the CFX96 Real-Time PCR detection system (BioRad) using the following parameters: 98 °C for 2 min followed by 39 cycles consisting of three steps – 98 °C for 5 sec, 60 °C for 5 sec and 72 °C for 5 sec with SYBR Green quantification at the end of each cycle. Melt curve analysis was conducted from 65 °C with a gradual increase in 0.5 °C increments every 0.05 sec to 95 °C with SYBR Green quantification conducted continuously throughout this stage. The raw data was analyzed using the Biorad CFX Manager 3.0 Software (BioRad).

#### Scanning electron microscopy (SEM) and transmission electron microscopy (TEM)

SEM and TEM were used to study the cell surface morphologies of the WT BCG and rBCG strains. The bacteria were immobilized to poly-l-lysine charged coverslips for 30 min and processed for SEM. Similarly, for TEM, bacterial suspensions were fixed and embedded in Spurr’s resin. The immobilized bacteria were rinsed with phosphate buffered saline (PBS), and fixed in 2.0% paraformaldehyde, 2.0% glutaraldehyde in 1X PBS with 3 mM MgCl2, pH 7.2 for 1 hour at room temperature. This was followed by 3 cycles of 10 minute washes in sodium cacodylate buffer with 3% sucrose, samples were post-fixed in 0.8% potassium ferrocyanide, 1% OsO_4_ and 3mM CaCl_2_ in 0.1 M sodium for 1 hour on ice in the dark. Samples were then rinsed in sodium cacodylate buffer and slowly rocked at 4ºC overnight. After a brief water rinse (2 × 5 min), bacteria were placed in 2% uranyl acetate for 1 hour at room temperature in the dark. The samples were dehydrated through a graded series of ethanol to 100% EtOH, then a 1:1 solution of ethanol:Hexamethyldisiloxazne (HMDS) (Polysciences) followed by pure HMDS. Coverslips were dried in a desiccator overnight and then attached to aluminum stubs via carbon sticky tabs (TedPella Inc.), and coated with 20 nm of AuPd with a Denton Vacuum Desk III sputter coater. Stubs were viewed and digital images captured on a Leo 1530 field emission SEM operating at 1 kV. For TEM, equal volumes of 2X fixative (as described above) were added to bacterial suspensions and rocked for 10 minutes at room temperature. Samples were centrifuged, supernatant removed and 1X fixative added carefully to not disturb the pellet. All subsequent steps were identical to the protocol described above up for SEM to the final 100% ethanol step. Bacterial cells were transferred to propylene oxide, and gradually infiltrated with Spurr’s low viscosity resin (Polysciences): propylene oxide. After 3 changes in 100% Spurr’s resin, pellets were cured at 60ºC for two days. Sections were cut on a Reichert Ultra cut E with a Diatome Diamond knife. 80 nm sections were picked up on formvar coated 1 × 2 mm copper slot grids and stained with tannic acid and uranyl acetate followed by lead citrate. Grids were viewed on a Phillips CM 120 TEM operating at 80 kV and digital images captured with an AMT 8 K x 8 K CCD camera.

#### Flow cytometry

Flow cytometry was used for analysis of fluorophore labeled cells. Cells were grown in 5 ml of Middlebrook 7H9 broth supplemented with appropriate antibiotics at 37ºC with shaking to an OD_600nm_ of 0.6. Thereafter, 1 ml of the culture was labelled with TetraFI (TAMRA-L-Ala-D-glutamine-L-Lys-D-Ala) for 3 hours at 37ºC with shaking. The CytoFLEX flow cytometer (Beckman Coulter) was used for analysis of TetraFI labeling.

#### Mammalian cell culture

For cell-based *ex vivo* infection assays, the human monocyte U937 cell line (obtained as a gift from the Council for Scientific and Industrial Research of South Africa [CSIR]) was grown in RPMI-Glutamax (Cat. 61870-036, Fischer Scientific) supplemented with 10% heat inactivated fetal bovine serum (FBS) (Cat. 10082147, Fischer Scientific) at 37ºC with 5% CO_2_. In addition, BMDMs extracted from the bone marrow (BM) of 6-8 weeks old female wildtype BALB/c mice were cultivated in a similar manner. BMDMs were generated as previously described by Toda et al. (Toda et al., 2021). Briefly, for differentiation of BM cells into macrophages, BM cells were seeded in BMDM differentiation media (RPMI-Glutamax supplemented with 10% FBS and 10% L929-conditioned media) and differentiated for 6 days. Non-adherent cells were washed out with warm BMDM differentiation media and adherent macrophages were used for *in vitro* infection assays.

#### Mtb containment following *in vitro* training with BCG, rBCG or other antigens in human monocytic U937 cell lines

*In vitro* training of monocytes was performed according to the model by Bekkering et al, 2016 (Bekkering et al., 2016) and Pan et al. (Pan et al., 2020). Briefly, U937 monocytes (1 × 10^6^/mL) were transferred into a 24-well plate and cells were incubated with either culture medium only as a negative control or MDP, LPS, heat killed WT BCG or heat killed rBCG::iE-DAP at 37 °C, and 5% CO_2_ for 24 h. Cells were washed twice with 1 mL of warm PBS and then incubated for 2 days in RPMI with 10% FBS and penicillin-streptomycin in the presence of 25 nM phorbol 12-myristate 13-acetate (PMA) (which can induce the differentiation of monocytes to macrophages). After washing twice with 1 mL of warm PBS, the differentiated macrophages were infected with Mtb H37Rv at MOI:1 and incubated for 24h. After 24 h, cells were lysed and bacterial load was enumerated by plating for CFU counts on 7H11 Middlebrook media.

#### Enzyme-linked immunosorbent assay (ELISA)

Sandwiched ELISA was performed for cytokine (TNF-α) measurement in culture supernatants. Culture supernatants were used immediately after harvest for ELISA. Sandwiched ELISA (R&D systems) was performed as per manufacturer’s recommendations.

#### BCG infection of BALB/c mice and CFU enumeration

To determine the lung bacillary burden of wild-type and rBCG::iE-DAP strains 6-8 weeks-old female BALB/c mice were infected using the aerosol route in a Glascol inhalation exposure system (Glasscol). The inoculum implanted in the lungs at day 1 (n = 3 mice per group) in female BALB/c mice was determined by plating the whole-lung homogenate on 7H11-selective plates containing carbenicillin (50 mg/ml), Trimethoprim (20 mg/ml), Polymyxin B (25 mg/ml) and Cycloheximide (10 mg/ml). Doxycycline was administered at determined doses for CRISPRi activation by daily oral gavage and following infection, mice lungs were harvested (n = 5 animals/group), homogenized in sterile PBS and plated on 7H11-selective plates at different dilutions. The 7H11-selective plates were incubated at 37°C and single colonies were enumerated after 4 weeks for the 10 days aerosol infection experiment, and also after 4 weeks for the 8 weeks aerosol infection experiment.

#### Mouse immunization and determination of protective efficacy against Mtb infection

To test the efficacy of rBCG::iE-DAP as a vaccine candidate, BALB/c mice (n = 10 per group) were immunized intradermally with 10^5^ colony-forming units (CFU)/100 µL of WT BCG or rBCG::iE-DAP strains. Mice were sham immunized with saline (n=10) and Dox was administered by daily oral gavage to the Saline+Dox (n=5), WT BCG+Dox (n=5) and the rBCG::iE-DAP+Dox (n=5) groups for 6 weeks. Mice were weighed every week to monitor the effect of Dox administration on the health of the mice. Mice were challenged with ∼100 CFU of Mtb H37Rv strain by the aerosol route 6 weeks post immunization in a Glasscol inhalation exposure system (Glasscol). Lungs and spleens from infected animals were harvested at week 4 and week 8 post Mtb infection for analysis of lung bacterial burden by plating the whole-lung homogenate on 7H11-selective plates containing carbenicillin (50 mg/ml), Trimethoprim (20 mg/ml), Polymyxin B (25 mg/ml) and Cycloheximide (10 mg/ml) and lung pathology was assessed after hematoxylin and eosin (H&E) staining.

#### Histopathology

Half of the left lung/mouse was cut and fixed in 10% neutral buffered formalin, paraffin embedded, sectioned, and H&E stained. Slides were digitally scanned (Aperio AT turbo scanner console version 102.0.7.5; Leica Biosystems, Vista, CA), transferred (Concentriq for Research version 2.2.4; Proscia, Philadelphia, PA), and visualized (Aperio ImageScope version 12.4.0.5043; Leica Biosystems Pathology Imaging, Buffalo Grove, IL). Histology images were analyzed with the Fiji software (ImageJ version 1.47n [NIH]).

## Acknowledgements

We thank the SAMRC, Wits University research office and the Fulbright Scholarship program for providing M.T.S with scholarships to pursue this work. We thank Jeremy Rock for donating CRISPRi plasmids and Peter Sander for the BCG Pasteur strain. We thank members of the CBTBR and Bishai Lab for helpful discussions and input on the manuscript. We are grateful for the assistance by the Berger Lab, Johns Hopkins School of Medicine for performing SEM and TEM and the Sidney Kimmel Comprehensive Cancer Center, Johns Hopkins School of Medicine for performing histology of lung samples.

## Author contributions

MTS and BDK conceived the project. MTS performed experiments, analyzed data and prepared the figures and manuscript. MP, KLO and AJA provided the TetraFl-1 PG probe. WRB and PU assisted with experiment design, mice vaccination and infection experiments and supervised the experiments.

## Data and materials availability

All data are available in the main text or the supplementary materials.

## Supplementary Material

Supplementary Figure S1-S4

