## Supplementary Information for "A modified BCG with depletion of enzymes associated with peptidoglycan amidation induces enhanced protection against tuberculosis in mice"

**Supplementary Figure 1**

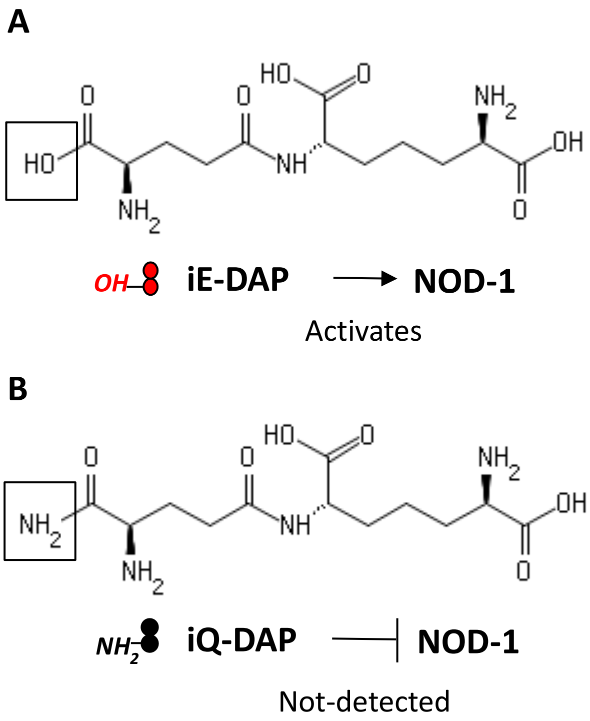


Supplementary Figure 1: **Molecular structures of iE-DAP and iQ-DAP**. iE-DAP is a NOD-1 ligand and modification of iE-DAP to iQ-DAP leads to evasion of NOD-1 activation.

**Supplementary Figure 2**

**A**

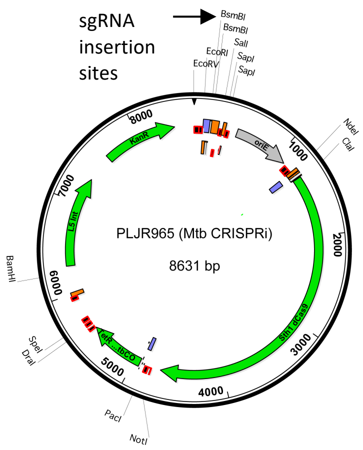


Plasmid PLRJ965 (Rock et al, 2017)

**B**

| rBCG::CRISPRi Strain | Gene Name | PAM Sequence | SgRNA Targeting Sequence (5'-3') |
|---|---|---|---|
| 1. rBCG::iE-DAP | Mb3739-Mb3740 **Operon** | TGAGCAA | AGCTGGTCTCGGGAGAGGT |

**C**

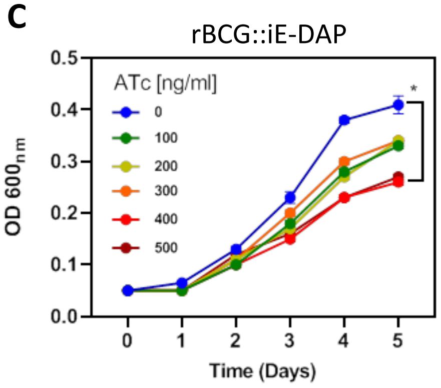


Supplementary Figure 2: **CRISPRi depletion of MurT-GatD in rBCG::iE-DAP**. (A) Plasmid PLRJ965 encoding dCas9 endonuclease from *S. thermophiles* and for expression of inserted sgRNA. (B) Table with gene targets and sgRNA targeting sequences. (C) Growth kinetics of rBCG::iE-DAP grown in a range of ATc [0-500 ng/ml]. rBCG::iE-DAP grown media without Atc grows at a similar rate as WT BCG. Activation of the CRISPRi platform in rBCG::iE-DAP with ATc [100-500 ng/ml] resulted in reduced growth in a concentration dependent manner.

### Supplementary Figure 3

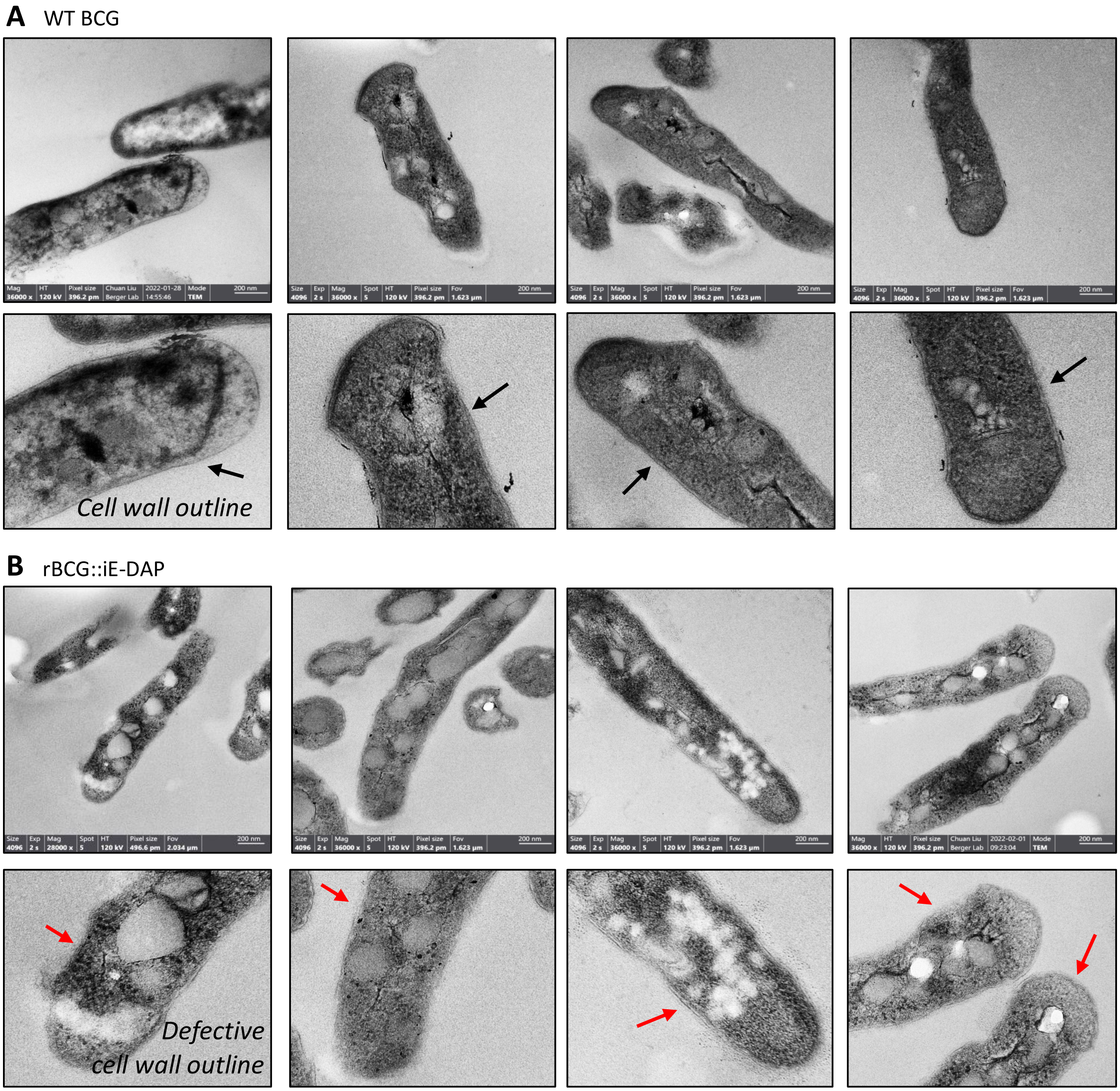


<u>Supplementary Figure 3:</u> **TEM reveals defective cell wall of rBCG::iE-DAP**. Transmission electron micrographs of WT BCG (A) and rBCG::iE-DAP (B) grown in media supplemented with 200 ng/ml ATc. Depletion of MurT-GatD causes cell wall defects. Black arrows denote clear cell wall structure. Red arrows denote defective cell wall.

**Supplementary Figure 4**

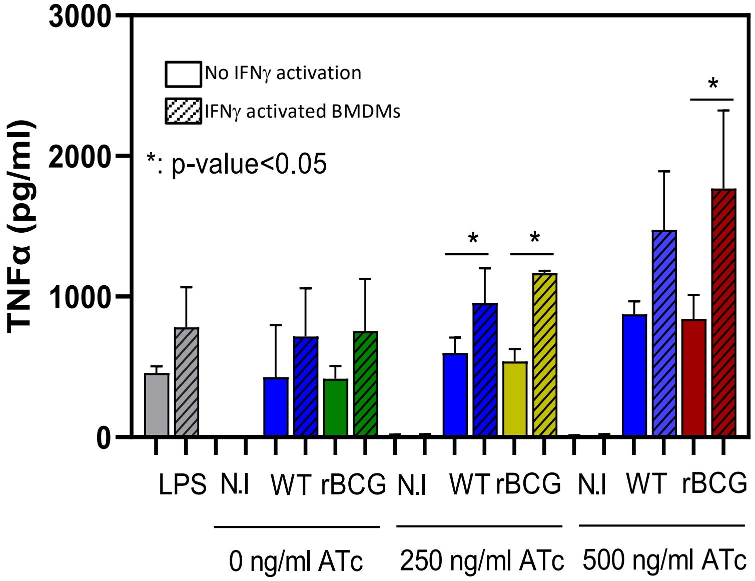


Supplementary Figure 4: **Analysis of secreted TNFα levels from non-activated and IFNγ-activated BMDMs infected with WT BCG and rBCG::iE-DAP at MOI 1:20.** Increased TNFα secretion was observed from rBCG::iE-DAP infected IFNγ-activated BMDMs cultured in media supplemented with 500 ng/ml ATc. LPS was used as a control.
